# A Transcriptomic Analysis of Xylan Mutants Does Not Support the Existence of A Secondary Cell Wall Integrity System in Arabidopsis

**DOI:** 10.1101/246330

**Authors:** Nuno Faria Blanc, Jenny C. Mortimer, Paul Dupree

## Abstract

Yeast have long been known to possess a cell wall integrity (CWI) system, and recently an analogous system has been described for the primary walls of plants (PCWI) that leads to changes in plant growth and cell wall composition. A similar system has been proposed to exist for secondary cell walls (SCWI). However, there is little data to support this. Here, we analysed the stem transcriptome of a set of cell wall biosynthetic mutants in order to investigate whether cell wall damage, in this case caused by aberrant xylan synthesis, activates a signalling cascade or changes in cell wall synthesis gene expression. Our data revealed remarkably few changes to the transcriptome. We hypothesise that this is because cells undergoing secondary cell wall thickening have entered a committed programme leading to cell death, and therefore a SCWI system would have limited impact. The absence of transcriptomic responses to secondary cell wall alterations may facilitate engineering of the secondary cell wall of plants.

## 1 Introduction

Cell walls are polysaccharide rich matrices which surround the cells of plants, fungi and bacteria. What makes plant cell walls exceptional is their importance to human society. We rely on plant cell walls for dietary fibre, animal feed, building materials, paper, and fuel, amongst other uses. Recent biotechnological advances, such as synthetic biology, combined with a detailed knowledge of cell wall biosynthesis, have raised the possibility of engineering an optimised plant secondary cell wall. For example, we and others have successfully engineered plants cell walls which have biomass designed for more efficient conversion to biofuels (Coleman et al., 2009;Mortimer et al., 2010;Petersen et al., 2012;Smith et al., 2013;Gondolf et al., 2014).

Unexpectedly, the engineered plants often grow normally, despite substantial biochemical and structural alterations, suggesting a tolerance for change.

Plant cell walls exist as two major types: primary and secondary. The primary wall is a thin layer that forms from the cell plate after mitosis. Since it surrounds growing cells, it must be flexible, enabling cell expansion either isotropically or anisotropically. Secondary cell walls are much thicker, can be lignified, and are laid down on the inside of the primary cell wall in some cell types, at the cessation of growth. The exact composition of these walls varies depending on cell function, tissue type and plant species. Arabidopsis has a primary cell wall composed of cellulose, xyloglucan and pectins, whereas its secondary cell wall is dominated by cellulose and xylan, with small amounts of pectin and mannan. Some secondary cell walls, such as those of the xylem water transport vessels, are extensively lignified after the deposition of the polysaccharides.

Due to the importance of the cell wall to an organism, it is perhaps unsurprising that cells might possess a system which detects and repairs damage whilst allowing the dynamic changes required of a growing cell. In yeast, this cell wall integrity system (CWI) has been well characterised. The yeast plasma membrane contains cell surface sensors which have a highly mannosylated serine threonine rich (STR) extracellular domain (Philip and Levin, 2001). These sensors are also thought to be anchored in the cell wall, with the STR acting as a nano-spring to detect stresses and strains between the wall and membrane (Kock et al., 2015). The signal transduction pathway is a mitogen-activated protein kinase (MAPK) cascade acting via the master regulator Rho1p, a small G protein (Levin, 2011). The various outputs include changes in polysaccharide biosynthesis and actin reorganisation.

The discovery of a cell wall integrity system in plant primary cell walls (PCWI) is much more recent, although it has been recognised for some time that perturbations to the primary cell wall can lead to compensatory effects. The activation of the PCWI system is linked to the process of control of growth and adaptation of cell wall architecture and composition to varying environments (Voxeur and Hofte 2016). Disruption of primary cell wall cellulose synthase activity is an activator of the PCWI. For example, the *ectopic lignification* (*eli1*) mutation of *CESA3* or application of the herbicide isoxaben, which targets cellulose biosynthesis, leads to ectopic lignification (Cano-Delgado et al., 2003). *prc1-1*, a mutation in *CESA6* also shows ectopic lignification and callose accumulation (Desprez et al., 2002;Hematy et al., 2007). Virus Induced Gene Silencing (VIGS) of a primary wall CESA in tobacco results in a compensatory increase in homogalacturonan (Burton et al., 2000).

Receptor like kinases (RLKs) have emerged as the likely mechanism by which signals are transduced into the plant cell. These proteins have an extracellular ligand-recognition domain, a transmembrane domain and a cytosolic kinase domain. Extracellular activation leads to phosphorylation of downstream signalling components. RLKs form a large gene family (more than 600 in Arabidopsis (Shiu and Bleecker, 2003)) and only a small proportion have been characterised in terms of ligand specificity and function. However, the roles described are extremely varied and include disease resistance, symbiosis, shoot apical meristem (SAM) maintenance and determination of cell fate. THESEUS1 (THE1), a plasma-membrane spanning Ser/Thr RLK, was first identified as a suppressor of the CESA6 mutant *prc1-1* (Hematy et al., 2007). THE1 was shown to mediate many, although not all, of the downstream outputs which result from inhibition of primary cell wall synthesis. The signal which activates RLKs such as THE1 is also unknown. Another plasma membrane spanning Ser/Thr RLK, WALL ASSOCIATED KINASE1 (WAK1), binds oligogalacturonides (OGs) (Decreux and Messiaen, 2005). OGs are oligosaccharide fragments of the pectin homogalacturonan, and are interpreted by the plant as DAMPs (damage associated molecular patterns). Detection of OGs by WAK1 leads to downstream activation of defence response genes and changes in growth and development (Brutus et al., 2010).

The evidence for a CWI system functioning in the plant secondary cell wall (SCWI) is much less apparent, although it has been widely proposed e.g. (Hamann, 2012; Doblin et al., 2014; Vermerris and Abril, 2014). There have been no clear reports that changes to the secondary cell wall composition induce compensatory changes in these tissues other than those simply due to developmental differences, for example due to dwarfing of the plant. Mutations in the secondary cell wall cellulose synthases (CESA4, 7 and 8), unlike the primary wall CESAs, do not result in widespread cell wall composition modifications beyond the directly caused reduction in cellulose. The *irx3* mutant (a point mutation in *CESA7*), has collapsed xylem vessels and less than a fifth of the cellulose content of wild type (WT) stems (Turner and Somerville, 1997), yet continues to make xylan and lignin. Solid-state NMR spectra of *irx3* and WT cell wall material revealed that the difference between the two samples was simply cellulose, indicating there were no observable changes to lignin or hemicellulose that would suggest adaptation to the loss of cellulose (Ha et al., 2002).

Plants which have reduced xylan synthase activity such as *irx9*, *irx10* or *irx14* have a secondary cell wall weakness sufficient to result in xylem vessel collapse, Nevertheless, they do not have large changes in wall composition (Brown et al., 2007). The alterations reported are consistent with their stunted development, and do not indicate a compensatory response. Further support for the view that the SCW is amenable to modification comes from the finding that the lignin composition is very plastic and can be altered through metabolic engineering without negative impact on the plant (Wilkerson et al., 2014; Eudes et al., 2015).

Perhaps the strongest evidence for a SCWI system comes from lignin biosynthesis mutants. Mutants in lignin biosynthesis which result in a reduction in lignin content, do show evidence of compensatory modifications to the cell wall. For example, silencing of *4-COUMARATE:COALIGASE1* (*4CL1*)/*4CL2* and *CAFFEOYL-COA O-METHYLTRANSFERASE* (*CCoAOMT*) in Arabidopsis and maize respectively results in an increase in cellulose content (Yang et al., 2011;Li et al., 2013). They show clear changes to transcription (Vanholme et al. 2012). However, this may be because lignification is not always a cell autonomous process. Parenchyma cells surrounding the xylem vessels act as “feeder cells”, providing the monolignol subunits (Petersen et al., 2012;Smith et al., 2013). Evidence for a detection system for a loss of lignification comes from recent work which used a similar approach to that used to identify THE1. A suppressor screen was performed on a dwarf mutant which has reduced *p-*coumaroylshikimate 3’-hydroxylase (3CH) activity, *reduced epidermal fluorescence8-1 (ref8-1)*, and it was shown that interference with two MEDIATOR (MED) complex subunits (MED5a and MED5b) rescues the growth phenotype of *ref8-1* without restoring lignin biosynthesis (Bonawitz et al., 2014). The MED complex is localised in the nucleus, so it is proposed that a currently unknown sensor detects the changed lignin precursor components, and relays that information, via the MED complex, to result in large scale transcriptional changes in the living cells surrounding the fibres and vessels (Bonawitz et al., 2014). Thus, the cells not undergoing secondary cell wall deposition may detect and respond to changes in phenylpropanoid intermediates.

Transcription factors (TFs) are key to regulating the development of the secondary cell wall. Since the decision to deposit secondary cell wall usually leads to cell death, plants have a cascade of TFs which control this process. These include VASCULAR-RELATED NAC DOMAIN6 (VND6) and VND7, which directly regulate the expression of the MYB46 TF and some secondary cell wall biosynthetic genes (Kubo et al., 2005;Zhong et al., 2008). A large-scale systems approach has recently identified that this feed-forward loop as one of many involved in regulating secondary cell wall biosynthesis (Taylor-Teeples et al., 2015). As yet, no TFs have been implicated in regulating a SCWI response, in which direction and balance are controlled, rather than turning on the cascade. However, if such a TF or network of TFs exists, we would predict that it is commonly upregulated in secondary cell wall synthesis mutants.

Xylan, the dominant secondary cell wall polysaccharide after cellulose, in composed of a (1-4)-β-D-xylose backbone, and depending on species and tissue, it is variously substituted with glucuronic acid, 4-*O*-methyl-glucuronic acid, arabinose and acetyl groups. In Arabidopsis, IRX9, IRX10 and IRX14 are the key proteins involved in secondary cell wall xylan backbone biosynthesis (Brown et al., 2007;Pena et al., 2007;Brown et al., 2009;Wu et al., 2010). Loss of function mutations in these genes lead to a reduced xylan chain length, although the severity is dependent on which gene is affected (Figure 1). In the absence of these genes, a reduced quantity of defective xylan is made by homologues IRX9L, IRX10L and IRX14L respectively (Brown et al., 2009;Wu et al., 2009;Wu et al. 2010. Arabidopsis secondary cell wall xylan is predominantly composed of (methyl)glucuronoxylan (Figure 1), where GLUCURONIC ACID SUBSITUTION OF XYLAN1 (GUX1) and GUX2 are responsible for glucuronic acid (GlcA) sidechain addition (Mortimer et al., 2010;Rennie et al., 2012), of which a proportion is subsequently methylated (Urbanowicz et al., 2012) (Figure 1). Since xylan biosynthesis is relatively well understood, we suggest that it provides a good system for in which to test whether Arabidopsis cells which are actively synthesising secondary cell wall, can detect and respond to defects in xylan biosynthesis.

**Figure 1:**
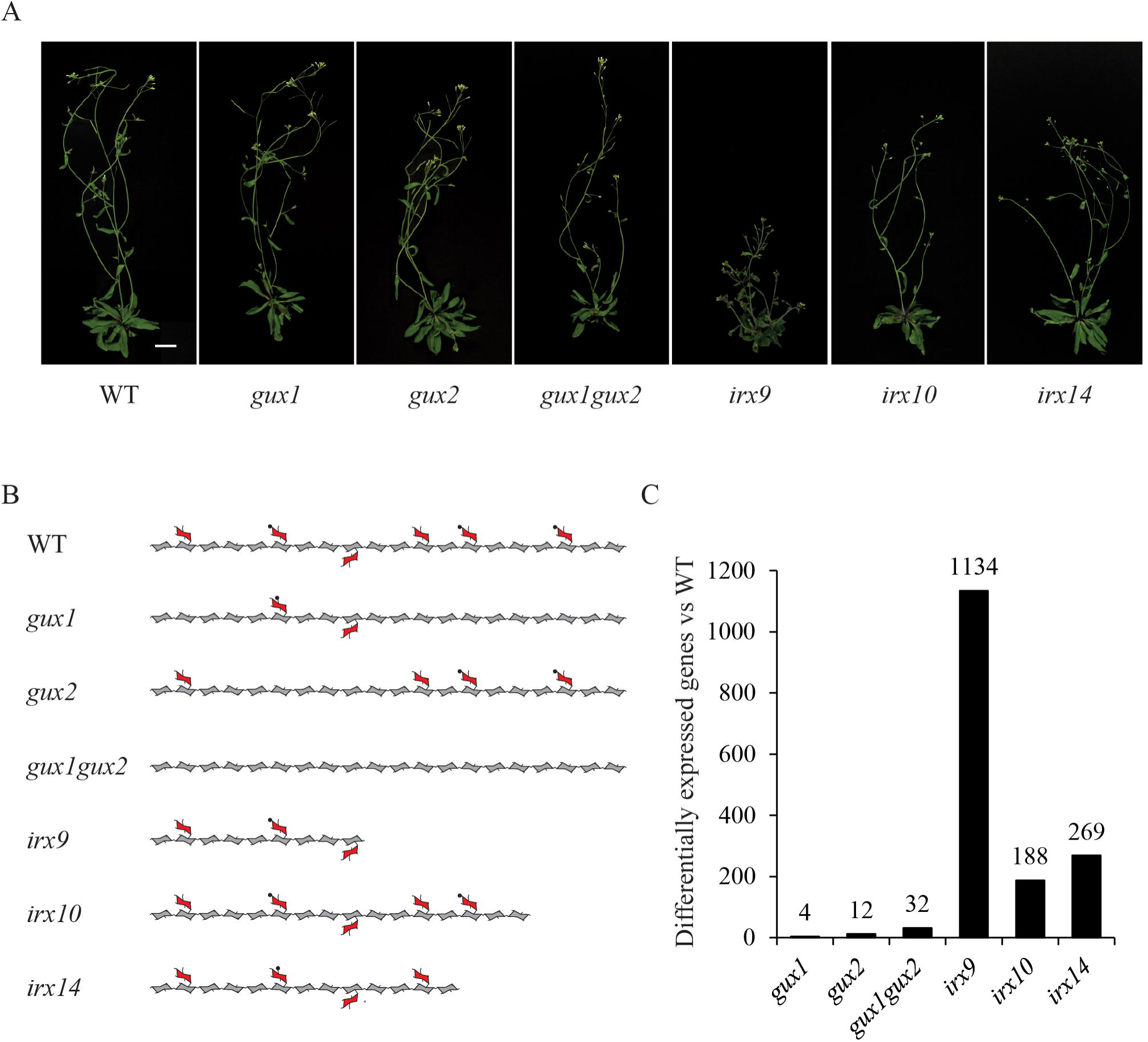
Xylan mutants used in this study. (**A**) Images of mutant plants used in this study. Scalebar = 20 mm. (**B**) Diagram representing the effects of these mutations on xylan structure. Acetylation and the reducing end oligosaccharide on the xylan molecule is not shown for clarity. Pentagons = xylose, hexagons = GlcA, triangles and circles = methyl groups. (**C**) Number of differentially expressed genes in each genotype compared to WT (fold change < 0.5 or > 2.0, *p*-value ≤ 0.05).

In this study, we used a transcriptomics approach to investigate whether we could detect putative components of a SCWI system in Arabidopsis stems. We looked at gene expression in a collection of xylan biosynthetic mutants. The aim of this experiment was threefold. Firstly, we wanted to test whether there is a co-ordinated expression response to xylan defects within the xylan biosynthetic pathway. Secondly, we wanted to identify more general responses to secondary cell wall defects, such as the upregulation of other biosynthetic pathways or transcription factors. Thirdly, we wanted to look for the upregulation of putative SCWI integrity sensors, such as RLKs.

## 2 Methods

### 2.1 Genotypes and growth conditions

The following previously published *Arabidopsis thaliana* T-DNA insertion lines were used: SALK_063763 (*gux1*, At3g18660), GABI-722F09 (*gux2*, At4g33330), SALK_057033 (*irx9*, At2g37090), SALK_046368 (*irx10*, At1g27440), SALK_038212 (*irx14*, At4g36890) and *gux1gux2*. All mutants were in the Columbia-0 (Col-0) ecotype. Seeds were stratified (4°C, 72 hours) prior to being grown on soil (Levington M3 compost) in a growth chamber (22°C, 60% humidity, continuous light). Several plants were initially grown in each pot but these were gradually thinned until only 2 plants remained to ensure all plants were of a similar developmental stage. All lines were genotyped by PCR prior to inclusion in this study, using the relevant published oligonucleotide primers.

### 2.2 Plant material

Plants were harvested at the same developmental stage (Boyes et al., 2001), defined as when the inflorescence stem containing an extended silique (1.5 cm) or presented silique fattening. At this point, inflorescence stem height was measured and basal stem with 1/5 of this height was harvested and immediately frozen in liquid nitrogen. Stems (15-20 per genotype) were pooled to form one biological replicate. Samples were harvested at the same time of day (within one hour) to reduce the effects of circadian differences on the data set. Three independently grown biological replicates were collected in this manner.

### 2.3 RNA extraction and microarrays

To extract RNA, tissue was ground under liquid N_2_ and mixed well with Trizol (1 mL per 100 mg of tissue; Invitrogen). Following incubation at room temperature for 2 minutes, chloroform (200 μL) was added and samples were gently mixed for 15 seconds. After centrifugation (12,000 × *g*, 4°C, 15 minutes), the aqueous upper phase was obtained to which 250 μL high salt solution (0.8 M sodium citrate, 1.2 M sodium chloride) and 250 μL isopropanol were added. After mixing and centrifuging the samples, the pellet was isolated and washed with 75% (v/v) and then 100% (v/v) ethanol. After resuspension, samples were incubated at 37°C for 30 minutes with DNase (Promega). Further sample processing was done using the RNeasy kit (Qiagen), according to the manufacturers “RNA Cleanup” protocol. RNA concentration and quality was measured using a BioPhotometer plus (Eppendorf) as well as by agarose gel electrophoresis. Samples were then sent to the Nottingham Arabidopsis Stock Centre (NASC; http://affymetrix.arabidopsis.info/) for analysis. Quality control was first performed using an Agilent Bioanalyzer to check the integrity of the RNA. Following preparation of cDNA, the samples were hybridized to the Arabidopsis ATH1 Genome Array (Affymetrix). Raw data can be obtained from NASCarrays, experiment number 668.

### 2.4 Microarray data analysis

Data analysis was done using the R-based FlexArray software (Blazejczyk et al., 2007). For the background correction of raw data, Gene Chip – Robust Multi-array Analysis (GC-RMA) was used (Wu and Irizarry, 2004). Arrays were normalised using quantile normalisation (Bolstad et al., 2003) and probeset summarisation used median polish (Berger and Carlon, 2011). Arrays were analysed for significance using the Significance Analysis of Microarrays test (Tusher et al., 2001), and probesets with a p-value ≤ 0.05 and at least 2-fold change of their log values were retained for further analysis. Gene annotations for the probesets were obtained from the Affymetrix website (http://www.affymetrix.com) and Gene Ontology (GO) was done using Classification Superviewer tool from the Bio-Analytic Resource for Plant Biology (http://bar.utoronto.ca) (Provart and Zhu, 2003). Data is presented as log_2_(abundance in mutant/abundance in WT), as described in (Vanholme et al., 2012).

## 3. Results and Discussion

### 3.1 Xylan mutants used in this study

The following Arabidopsis xylan mutants: *irx9*, *irx10*, *irx14*, *gux1*, *gux2* and *gux1gux2*, along with WT ecotype Col-0 plants (Table 1) were grown alongside each other until they developed their first silique. At this point, basal stems were harvested, and the RNA extracted for microarray analysis. Genotype verification was also performed, and all lines were shown by PCR to be homozygous for the expected T-DNA insertion (Supplemental Figure S1). The effect of each of these mutations on xylan structure, along with their phenotype at the developmental stage used in this study, is shown in Figure 1.

**Table 1:**
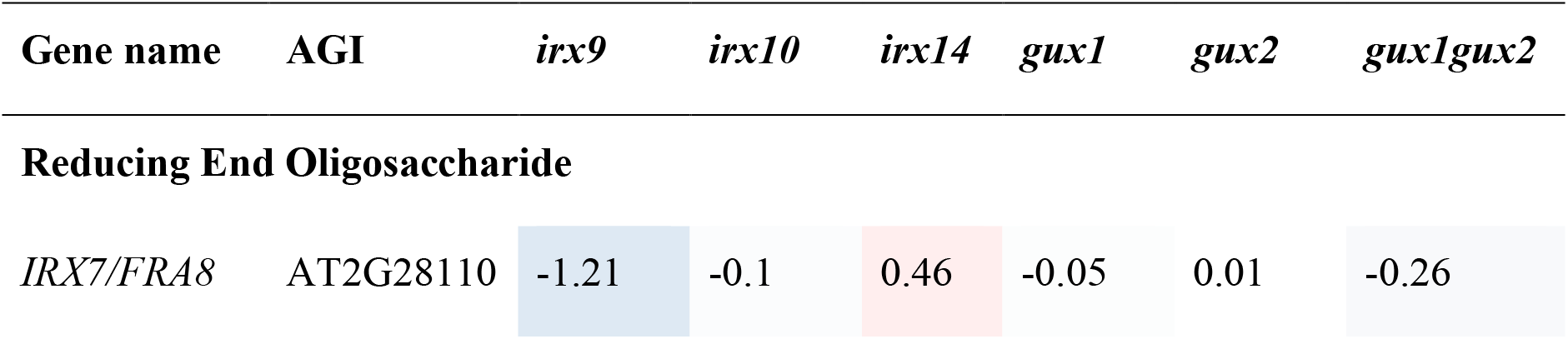
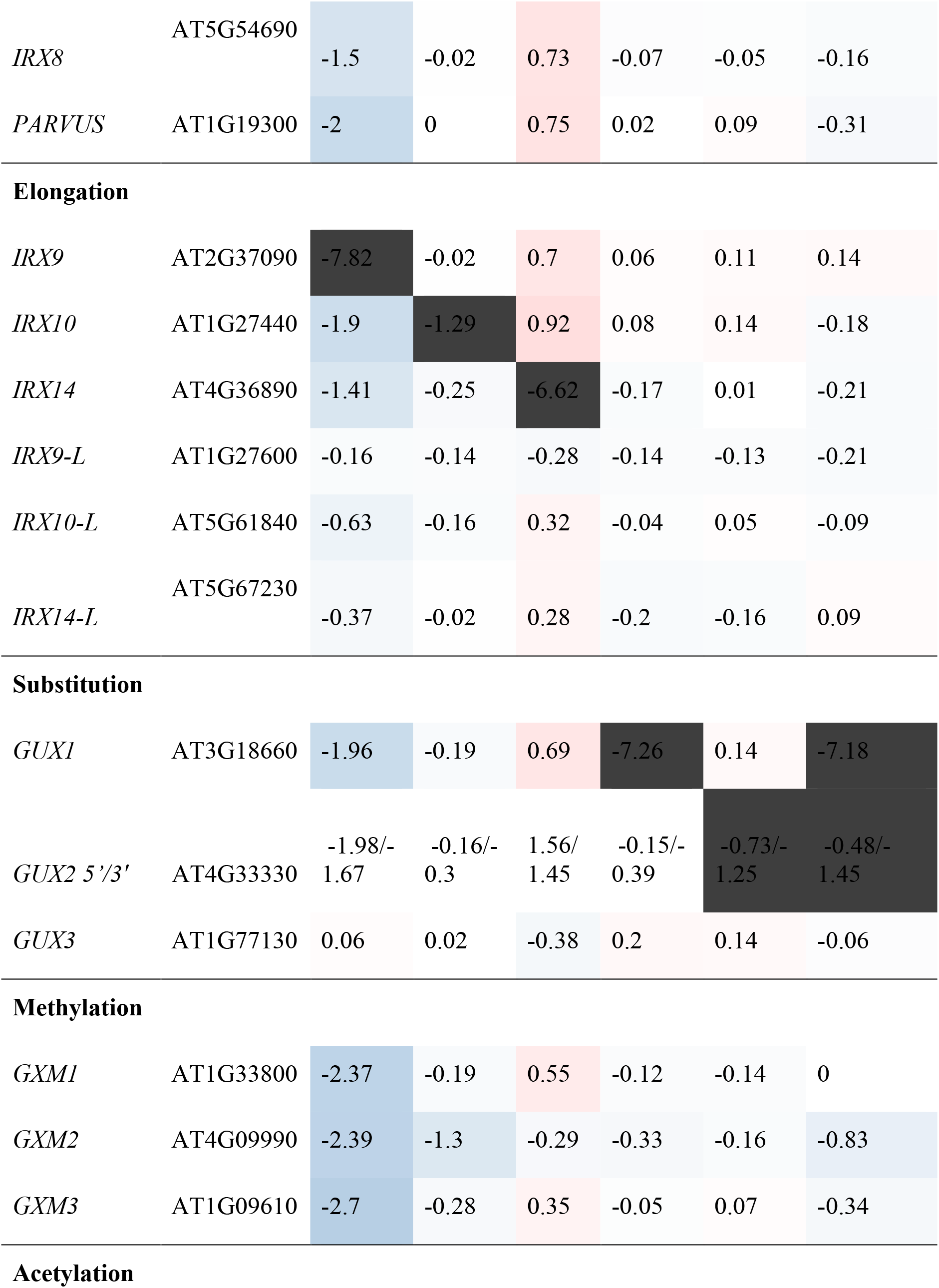
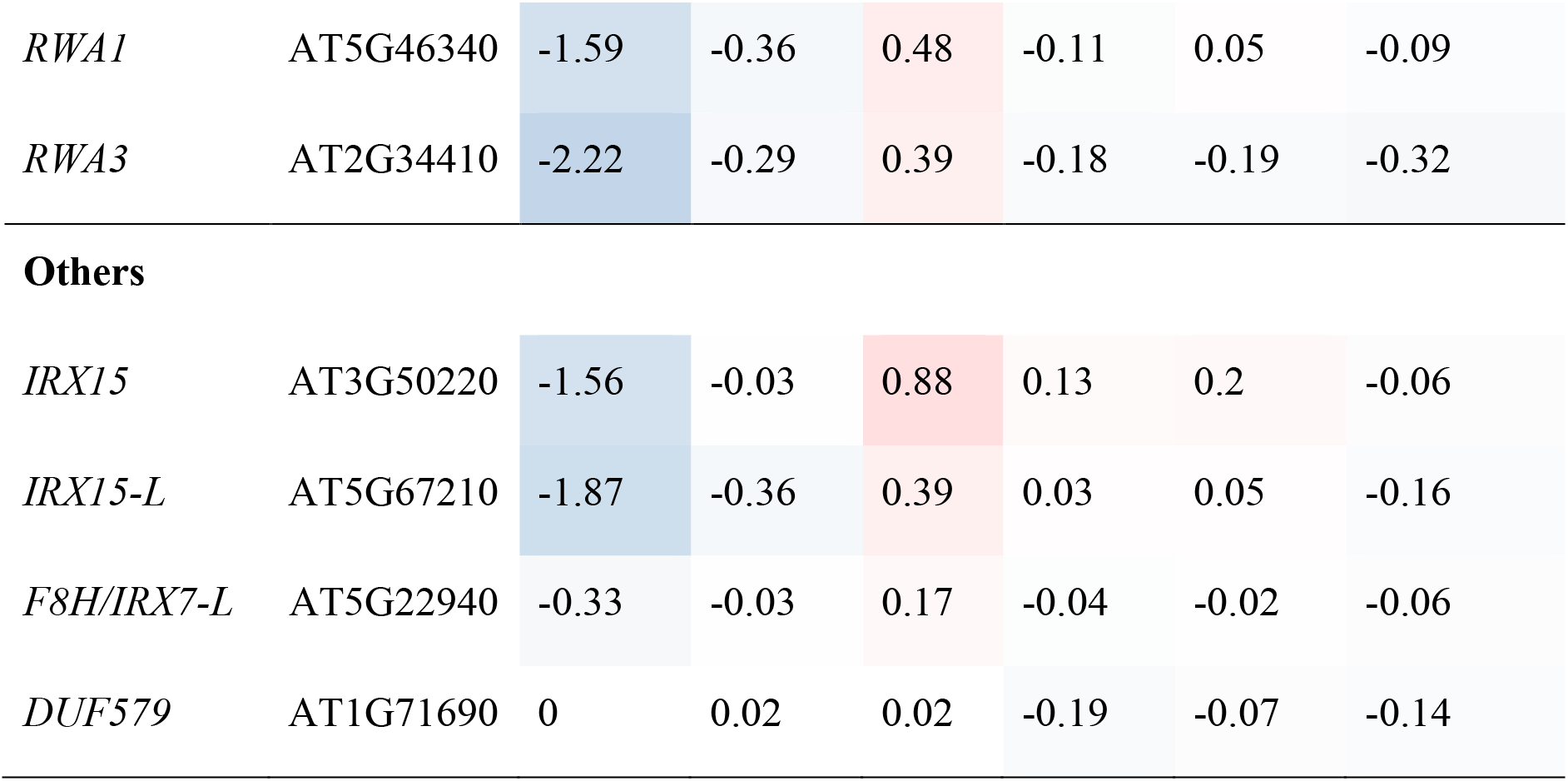
Expression of xylan synthesis-related genes represented on the Affymetrix ATH1 genechip in relation to WT. Values are shown as log_2_(abundance in mutant/abundance in WT). Values have been shaded red to indicate increased expression with respect to WT and blue to indicate decreased expression. Values in bold represent genes significantly different from WT. Grey shaded boxes indicate the mutant gene. GUX2 has two values corresponding to two probesets (earlier genome annotations assigned this region to two separate ORFs).

The microarray data were normalised (Supplemental Dataset 1) and statistically analysed, as described in the methods, to produce a set of genes which are up- or down-regulated in comparison to WT (Supplemental Dataset 2). The number of genes with a significant change relative to WT was surprisingly small for most of the mutants, and only *irx9* had more than a 1.5% change in its transcriptome. The extent of the change essentially followed the severity of the effect of the mutation on the plants gross morphology and development (Figure 1).

We confirmed that the tissue selected for analysis was still actively transcribing secondary cell wall synthesis genes. Expression data revealed that in the WT samples, genes associated with secondary cell wall biosynthesis, such as CESA4/IRX5 and IRX9, were all highly expressed (Supplemental Dataset 1).

### 3.2 Analysis of xylan-related gene expression

The microarray data confirmed that, for *irx9*, *irx14*, and *gux1* plants, the corresponding mutant gene had the largest decrease relative to WT (Table 1). However, for *irx10* and *gux2*, whilst transcript levels were reduced, they were still clearly present. This is likely due to the production of non-functional transcripts which can still bind to some of the individual probes in a probeset. For example, *irx10* (SALK_046368) has a T-DNA insertion in the middle of the fourth and final exon. Extraction of the raw probe data (Supplemental Dataset 1) reveals only partially reduced intensity for 5’ localised probes, but greatly reduced intensity for 3’ localised probes. The GUX2 open reading frame (ORF) was originally annotated as two different genes, and hence two different probesets can bind to it (here annotated 3’ and 5’ according to their location within the cDNA).

Next, expression of other xylan-synthesis related genes was explored to test whether a xylan synthesis feedback loop could be detected. For example, we hypothesised that if one gene involved in xylan backbone extension had reduced expression e.g. *IRX9*, resulting in reduced enzyme activity, then a close homologue might show compensatory increased expression e.g. *IRX9L*. Alternatively, if xylan elongation was affected, this could lead to an associated repression of downstream biosynthetic steps, such as GUX or glucuronoxylan methyl transferase (GXMT) activity, which might be apparent at the transcriptional level. Neither of these outcomes was visible in the data (Table 1). In *gux1*, *gux2*, *gux1gux2* and *irx10* essentially no significant differences in transcript levels from each other and from WT were observed (Table 1). Specifically, close homologues of the mutant genes did not show compensatory increases of expression e.g. *IRX10L* in *irx10* or *GUX2* in *gux1*. Whilst *irx9* showed a trend towards slight (but not significant) down regulation of xylan synthetic genes, and *irx14* showed a general (but again, not significant) trend for a small increase, we inferred that the plants were likely not exactly developmentally matched. Indeed, the xylan synthesis pathway is co-ordinately regulated with development of the stem (Brown et al., 2005). Since *irx9* and *irx14* had the largest gross morphological difference compared to WT (Figure 1A), matching the harvest developmental stage was the most challenging for these two genotypes. Overall, these data indicate that loss of xylan synthase activity does not result in altered gene expression of other xylan-related genes.

One alternative metabolic strategy to compensate for reduced quantities of enzyme is to boost the substrate pool to increase the rate of activity. Therefore, we analysed expression patterns of genes which have a role in synthesizing the UDP-sugars required for xylan biosynthesis. However, this subset of genes (Supplemental Dataset 3) revealed a very similar pattern to that reported for the xylan genes described above: a general small but significant decrease in *irx9*, and small but significant increase in *irx14*. Other mutants were essentially unaffected.

### 3.3 Pathways which provide substrates for non-sugar decorations of xylan show evidence of feedback in response to the reduced product sink

Xylan has various non-sugar modifications, including acetylation and methylation which impact the functional properties of the molecule. Mutants with short xylan chains (e.g. *irx9*, *irx10*, *irx14, irx15*) have an increased proportion of methylated (MeGlcA) versus unmethylated GlcA decorations, although the combined frequency of GlcA/MeGlcA ([Me]GlcA) substitution along the xylan backbone is maintained (Brown et al., 2007;Brown et al., 2011). This could be due to an active up-regulation of methylation activity to compensate for the shorter xylan chains, by altering the association between xylan and the amphiphilic surface of the lignin (Urbanowicz et al., 2014). Alternatively, it could be a passive effect resulting from the reduced pool of GlcA for the GXMT enzymes to act upon (Brown et al., 2007). Therefore, the expression patterns of the S-adenosylmethionine (SAM) cycle, the pathway responsible for recycling the SAM used for xylan GlcA methylation, and Acetyl-CoA (the substrate for TBL29 and other acetyl transferases) synthesis genes were investigated (Supplemental Dataset 3).

It has been previously suggested that the *gxmt* mutants accumulate SAM (Urbanowicz et al., 2012), and therefore we might predict that the *gux* mutants would also accumulate SAM. In fact, many of the SAM synthetases show reduced expression in *irx9* and *gux1gux2* implying that this pathway may respond transcriptionally to the reduced requirement for SAM in the Golgi. Since SAM is also the major methyl-donor for key metabolites, such as the hormone ethylene, vitamin B1 and polyamines (Hesse and Hoefgen, 2003), it is perhaps not surprising that levels are tightly controlled.

In the *gux* mutants, which lack GlcA, there is an increased amount of mono-acetylation (Chong et al., 2014) and the growth suppression seen in the xylan acetyl transferase mutant, *tbl29*, can be rescued by overexpression of GUX1 (Xiong et al., 2015), suggesting that increases in levels of xylan decoration could be beneficial to the cell wall. Acetyl-CoA, the precursor for xylan acetylation is synthesised via 2 independent pathways: from pyruvate in the mitochondria and plastid, and from citrate in the cytosol. The acetyl Co-A is then transported from the cytosol into the Golgi, likely by the RWA proteins (Manabe et al., 2011), where it is proposed to acetylate an intermediate (potentially AXY9 (Schultink et al., 2015)) prior to transfer of the acetyl group to the xylan backbone by the TBL29 acetyltransferase (Xiong et al., 2013;Urbanowicz et al., 2014). In our data, the cytosolic pathway was strongly downregulated, particularly in *irx9*, implying that acetyl Co-A synthesized from citrate is the major source of substrate for xylan acetylation, and that the size of the pool can be detected and used to regulate the cytosolic pool size.

### 3.4 Biosynthetic genes for non-xylan cell wall polysaccharides are unaffected

If xylan is altered in the plant, we may predict that an alternative hemicellulose such as mannan would be upregulated to compensate. Indeed, in gymnosperms mannan is the dominant hemiceullose, implying that xylan and mannan could be functionally interchangeable. Previously published biochemical analyses of these xylan mutants have shown that major changes to the cell wall are limited to the xylan itself (Brown et al., 2007;Brown et al., 2009;Mortimer et al., 2010), although it is possible that minor changes were not visible in these analyses. However, in agreement with this, transcriptional data showed no significant changes to genes involved in cellulose, mannan, xyloglucan or pectin synthesis, other than in the pattern recorded above that we have ascribed to developmental issues (decrease in *irx9*, small increase in *irx14*) (Supplemental Dataset 3). Taken together, these results suggest that defects in xylan synthesis do not cause major changes to the expression of genes that are involved in polysaccharide biosynthesis.

### 3.5 Alternative lignin biosynthetic genes are upregulated in the xylan mutants

Lignin has previously been shown to be deposited in primary cell walls in response to stress, including in cell wall synthesis mutants (Desprez et al., 2002;Cano-Delgado et al., 2003;Hematy et al., 2007), but not in secondary cell wall synthesis mutants. The exception to this is lignin mutants. Mutations in some lignin biosynthetic genes can result in compensatory upregulation of other parts of the lignin biosynthetic pathway, as well as increases in some matrix polysaccharides, likely pectins (Van Acker et al., 2013). A systems approach to understanding phenolic metabolism has revealed how lignin biosynthesis responds to perturbations (Vanholme et al., 2012). At least eleven enzymes, encoded by multiple genes, have been implicated in monolignol biosynthesis, although only a subset are associated with stem lignin biosynthesis (Table 2, marked in bold (Costa et al., 2003;Goujon et al., 2003;Vanholme et al., 2012;Vanholme et al., 2013)).

Here, the gene expression analysis in the xylan synthesis mutants (Table 2) showed trends that appear distinct from that seen for other groups of cell-wall related genes (Supplemental Dataset 3). In *irx9*, most of the lignin-related genes were significantly down-regulated (Table 2), which is supported by data showing that *irx9* has an overall decrease in lignin (Petersen et al., 2012). *PAL3*, *CCoAOMT3*, *CAD1* and *CAD3* are the exceptions to this, and were significantly upregulated. None of these isoforms are the dominant stem isoforms, implying that a different subset of the lignin pathway may be activated. The other xylan mutants showed milder alterations, with a general trend towards down-regulation of lignin-related genes, except for *CAD3* in *irx10* and *CCoAMT2* in *irx14*, which were upregulated.

**Table 2:**
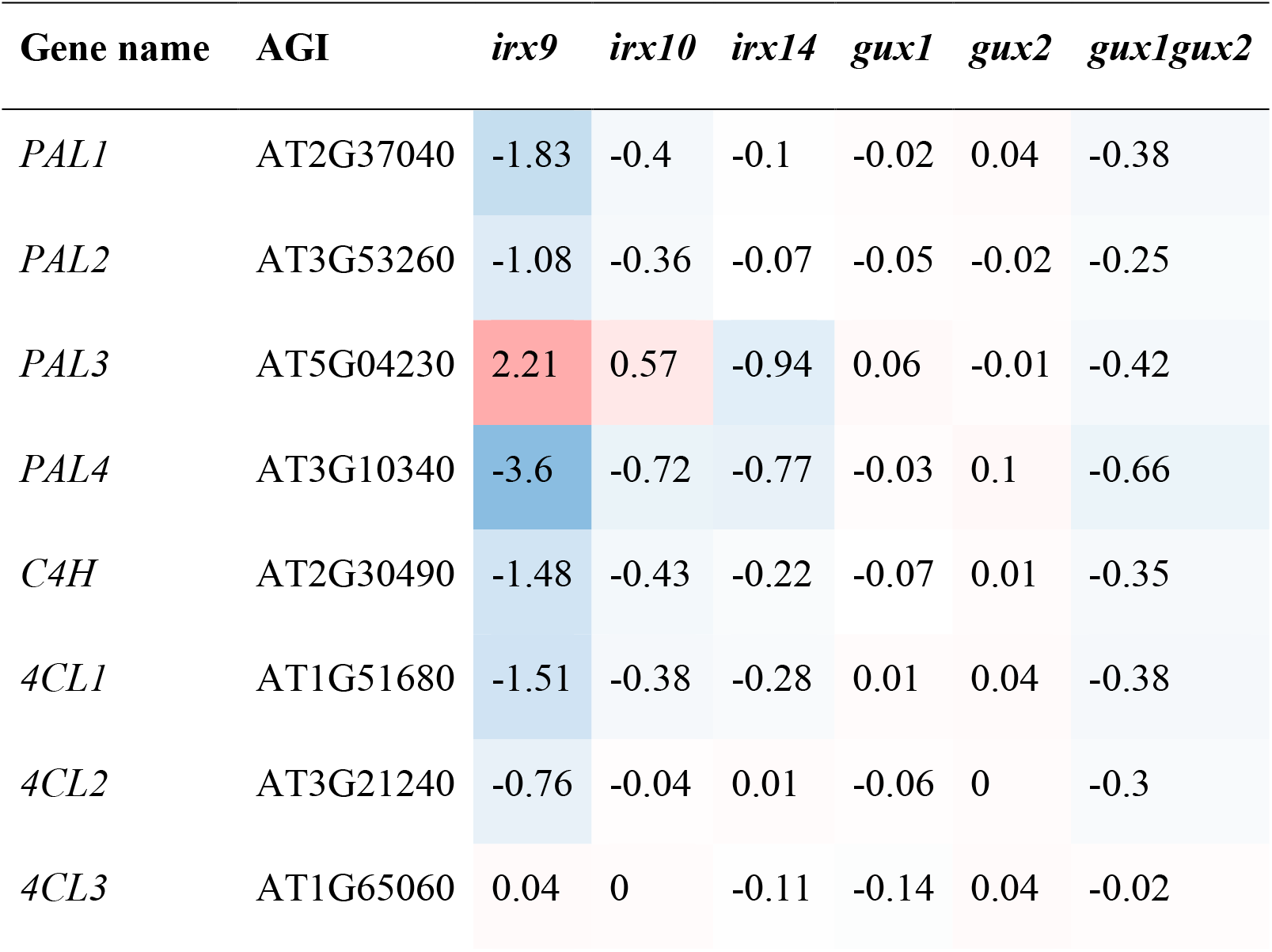
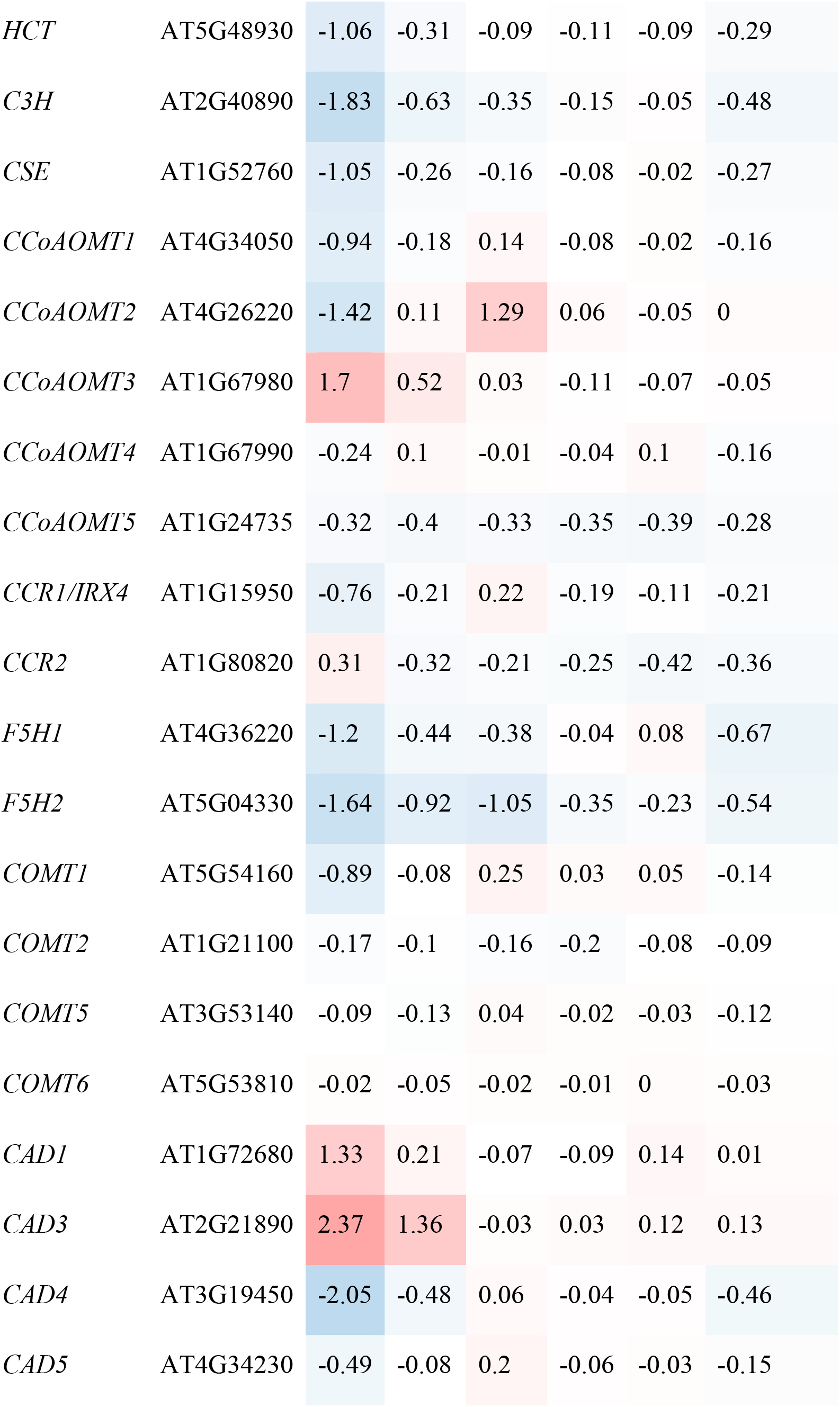
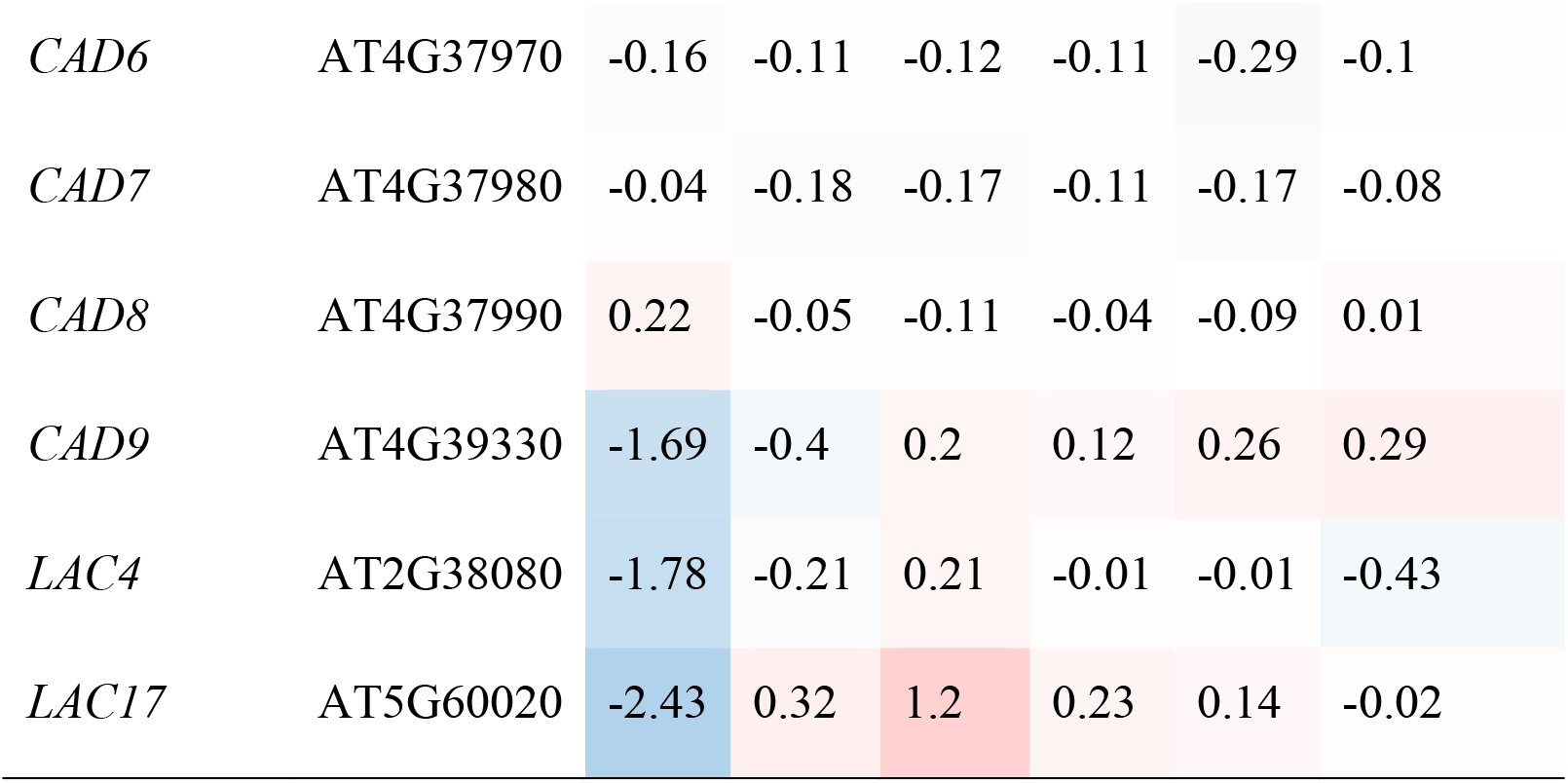
Expression of lignin biosynthetic genes represented on the ATH1 chip in relation toWT. Values are shown as log_2_(abundance in mutant/abundance in WT). Values have been shaded red to indicate increased expression with respect to WT and blue to indicate decreased expression. Values in bold represent genes significantly different from WT. Gene names/AGI in bold represent genes which are responsible for the majority of stem lignin biosynthesis.

| Gene name | AGI | *irx9* | *irx10* | *irx14* | *gux1* | *gux2* | *gux1gux2* |
| --- | --- | --- | --- | --- | --- | --- | --- |
| *PAL1* | AT2G37040 | -1.83 | -0.4 | -0.1 | -0.02 | 0.04 | -0.38 |
| *PAL2* | AT3G53260 | -1.08 | -0.36 | -0.07 | -0.05 | -0.02 | -0.25 |
| *PAL3* | AT5G04230 | 2.21 | 0.57 | -0.94 | 0.06 | -0.01 | -0.42 |
| *PAL4* | AT3G10340 | -3.6 | -0.72 | -0.77 | -0.03 | 0.1 | -0.66 |
| *C4H* | AT2G30490 | -1.48 | -0.43 | -0.22 | -0.07 | 0.01 | -0.35 |
| *4CL1* | AT1G51680 | -1.51 | -0.38 | -0.28 | 0.01 | 0.04 | -0.38 |
| *4CL2* | AT3G21240 | -0.76 | -0.04 | 0.01 | -0.06 | 0 | -0.3 |
| *4CL3* | AT1G65060 | 0.04 | 0 | -0.11 | -0.14 | 0.04 | -0.02 |
| *HCT* | AT5G48930 | -1.06 | -0.31 | -0.09 | -0.11 | -0.09 | -0.29 |
| *C3H* | AT2G40890 | -1.83 | -0.63 | -0.35 | -0.15 | -0.05 | -0.48 |
| *CSE* | AT1G52760 | -1.05 | -0.26 | -0.16 | -0.08 | -0.02 | -0.27 |
| *CCoAOMT1* | AT4G34050 | -0.94 | -0.18 | 0.14 | -0.08 | -0.02 | -0.16 |
| *CCoAOMT2* | AT4G26220 | -1.42 | 0.11 | 1.29 | 0.06 | -0.05 | 0 |
| *CCoAOMT3* | AT1G67980 | 1.7 | 0.52 | 0.03 | -0.11 | -0.07 | -0.05 |
| *CCoAOMT4* | AT1G67990 | -0.24 | 0.1 | -0.01 | -0.04 | 0.1 | -0.16 |
| *CCoAOMT5* | AT1G24735 | -0.32 | -0.4 | -0.33 | -0.35 | -0.39 | -0.28 |
| *CCR1/IRX4* | AT1G15950 | -0.76 | -0.21 | 0.22 | -0.19 | -0.11 | -0.21 |
| *CCR2* | AT1G80820 | 0.31 | -0.32 | -0.21 | -0.25 | -0.42 | -0.36 |
| *F5H1* | AT4G36220 | -1.2 | -0.44 | -0.38 | -0.04 | 0.08 | -0.67 |
| *F5H2* | AT5G04330 | -1.64 | -0.92 | -1.05 | -0.35 | -0.23 | -0.54 |
| *COMT1* | AT5G54160 | -0.89 | -0.08 | 0.25 | 0.03 | 0.05 | -0.14 |
| *COMT2* | AT1G21100 | -0.17 | -0.1 | -0.16 | -0.2 | -0.08 | -0.09 |
| *COMT5* | AT3G53140 | -0.09 | -0.13 | 0.04 | -0.02 | -0.03 | -0.12 |
| *COMT6* | AT5G53810 | -0.02 | -0.05 | -0.02 | -0.01 | 0 | -0.03 |
| *CAD1* | AT1G72680 | 1.33 | 0.21 | -0.07 | -0.09 | 0.14 | 0.01 |
| *CAD3* | AT2G21890 | 2.37 | 1.36 | -0.03 | 0.03 | 0.12 | 0.13 |
| *CAD4* | AT3G19450 | -2.05 | -0.48 | 0.06 | -0.04 | -0.05 | -0.46 |
| *CAD5* | AT4G34230 | -0.49 | -0.08 | 0.2 | -0.06 | -0.03 | -0.15 |
| *CAD6* | AT4G37970 | -0.16 | -0.11 | -0.12 | -0.11 | -0.29 | -0.1 |
| *CAD7* | AT4G37980 | -0.04 | -0.18 | -0.17 | -0.11 | -0.17 | -0.08 |
| *CAD8* | AT4G37990 | 0.22 | -0.05 | -0.11 | -0.04 | -0.09 | 0.01 |
| *CAD9* | AT4G39330 | -1.69 | -0.4 | 0.2 | 0.12 | 0.26 | 0.29 |
| *LAC4* | AT2G38080 | -1.78 | -0.21 | 0.21 | -0.01 | -0.01 | -0.43 |
| *LAC17* | AT5G60020 | -2.43 | 0.32 | 1.2 | 0.23 | 0.14 | -0.02 |

A potential answer to why lignin deposition seems to be the exception in terms of responsive 2CW modifications comes from recent papers which have explored the spatial and temporal nature of lignin deposition in Arabidopsis (Pesquet et al., 2013;Smith et al., 2013) and poplar (Gorzsás et al., 2011). These papers have provided strong evidence for the “good neighbour hypothesis”, in which it has been proposed that neighbouring, non-lignifed cells monolignols to lignifying cells (Hosokawa et al., 2001), at least in some cell types. This evolving view of lignification was recently reviewed by Voxeur and colleagues who proposed two models of lignification: cell autonomous lignification (CAL) and non-cell autonomous lignification (NCAL). In their view of cell wall lignification, the role of CAL and NCAL will depend of cell maturation and type. For example, fibre cells are predominantly lignified by CAL and supported by NCAL, whereas vessels are initially lignified by CAL prior to PCD but NCAL is then key to completion of the lignification process (Voxeur et al., 2015).

### 3.6 Known secondary cell wall associated transcription factors do not respond to changes in xylan content or structure

Expression changes in secondary-wall related transcription factors (TFs) were also evaluated (Supplemental Dataset 3). The pattern obtained was like that we have shown for multiple other sets of genes i.e. in *irx9*, most of these TFs had reduced expression compared to WT, whilst in *irx14* many were slightly but significantly upregulated. Few changes were seen in the other mutants. Importantly, we did not see the strong upregulation of a TF or network of TFs that might form a SCWI response network (Figure 2).

**Figure 2:**
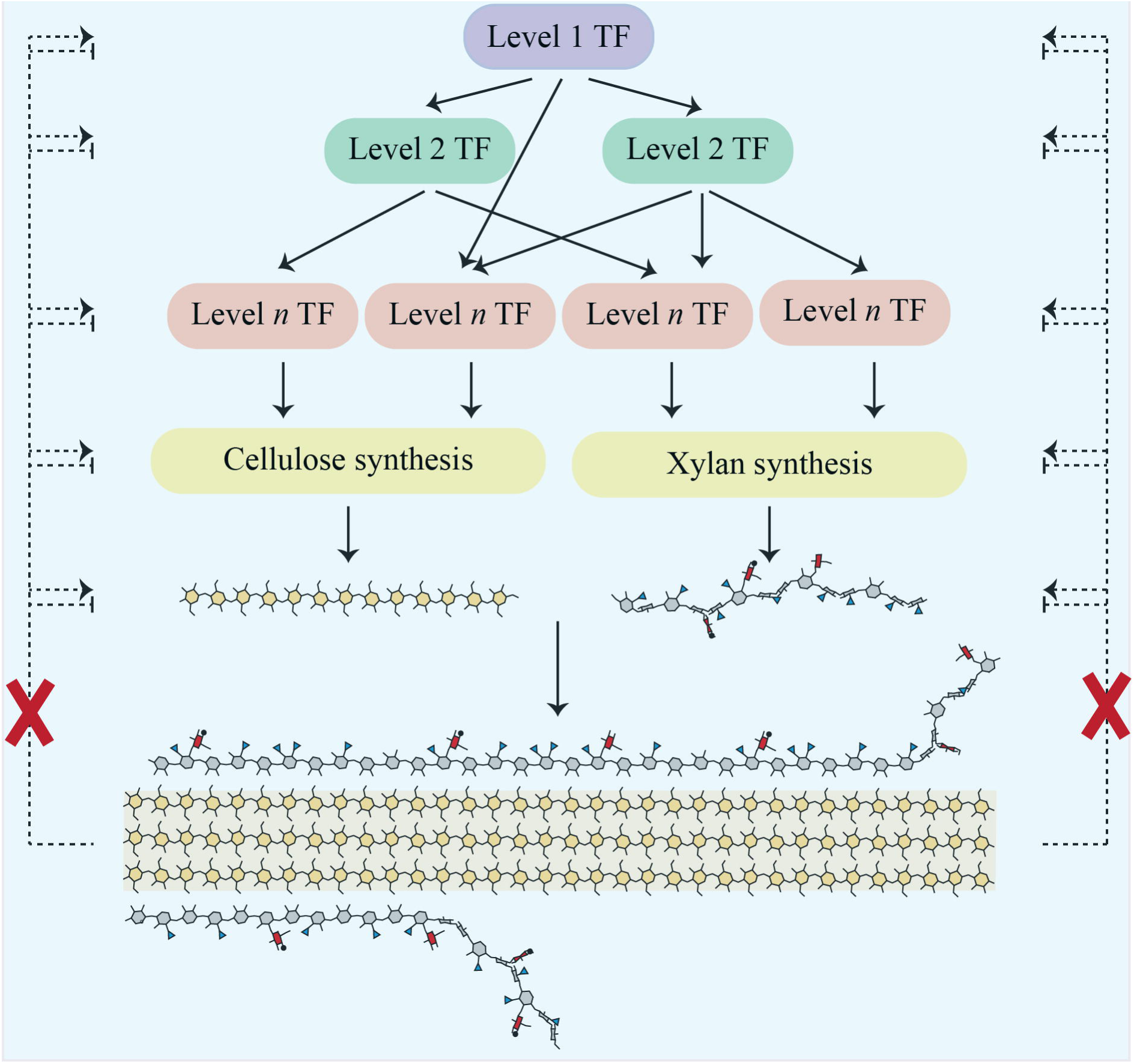
Scheme showing a cascade of transcription factors responsible for regulating cellulose and xylan biosynthesis in Arabidopsis secondary cell walls. Proposed regulatory feedback pathways which would induce a SCWI response are shown as dotted lines. Our work does not support their existence, as indicated by the red crosses.

### 3.7 Receptor Like Kinases (RLKs) transcripts do not show a consistent response to secondary cell wall changes

As described in the introduction, RLKs are the most likely candidates for detection of a loss of cell wall integrity, although due to their number (~600 in Arabidopsis), they have proven to be difficult to study. Data was extracted for all those RLKs on the ATH1 microarray (Supplemental Dataset 3) and searched for RLKs which showed a significant change in expression in at least two of the xylan mutants. However, none of the genes matched these criteria, including those associated with the PCWI pathways (Table 3). It should be noted that unchanged RLK expression does not exclude the possibility of RLK activation, as is seen for PCWI RLKs, such as THE1.

**Table 3:** Expression of RLK and other genes implicated in the PCWI response in relation to WT.Values are shown as log_2_(abundance in mutant/abundance in WT). Values have been shaded red to indicate increased expression with respect to WT and blue to indicate decreased expression. Values in bold represent genes significantly different from WT. Note that BZR1 and BZR2 bind to the same set of oligonucleotide probes.

| Gene name | AGI | *irx9* | *irx10* | *irx14* | *gux1* | *gux2* | *gux1gux2* |
| --- | --- | --- | --- | --- | --- | --- | --- |
| *BAK1* | AT4G33430 | -0.07 | -0.10 | 0.052 | 0.024 | 0.12 | 0.016 |
| *BRI1* | AT4G39400 | -0.48 | -0.20 | -0.09 | -0.03 | -0.08 | -0.09 |
| *HERK1* | AT3G46290 | -0.27 | 0.01 | -0.08 | 0.09 | 0.00 | 0.00 |
| *HERK2* | AT1G30570 | -0.11 | 0.10 | 0.00 | 0.16 | 0.14 | 0.07 |
| *THE* | AT5G54380 | -0.35 | -0.07 | 0.20 | -0.01 | -0.16 | -0.27 |
| *FER* | AT3G51550 | 0.19 | 0.09 | 0.07 | 0.00 | -0.02 | -0.10 |
| *MCA1* | AT4G35920 | **-1.32** | -0.13 | 0.33 | -0.16 | -0.08 | -0.16 |
| *BIN2* | AT4G18710 | 0.02 | 0.04 | -0.06 | -0.10 | -0.05 | -0.14 |
| *BZR1/BZR2* | AT1G75080/<br>AT1G19350 | -0.46 | -0.22 | -0.11 | -0.11 | -0.15 | -0.05 |

### 3.8 Abiotic and biotic stress associated genes are upregulated

There has been limited research on the wider effect of secondary cell wall modifications on plant physiology. Since changes to xylan structure, when severe enough, have a profound effect on plant growth (as in *irx9*), a more general analysis was performed on the transcriptomics data. The aim was to identify other signalling and response pathways which respond to gross cell wall defects. For example, the shorter xylan chains in *irx9*, *irx10* and *irx14* result, to varying extents, in collapsed xylem vessels. This limits water transport and is likely to induce drought stress, which we predict would be detected by abiotic stress response pathways. It was also recently shown that when plants are exposed to severe abiotic stress, expression of specific secondary cell wall TFs are enhanced. For example, under salt stress, VND7 expression expands to non-stele cells in the root, resulting in an extra strand of metaxylem (Taylor-Teeples et al., 2015).

In line with our prediction, gene ontology (GO) analysis of genes upregulated in *irx9* revealed an over-representation of genes associated with abiotic and biotic stress responses (Supplemental Figure S2). This pattern was repeated, although to a lesser extent, in the other *irx* mutants, but not in the *gux* mutants, as might be expected because they do not show collapsed xylem (Supplemental Figure S2). Further experiments are now required to exclude whether the stunting of *irx9* is due to the reduced xylan or poor water transport, or as we consider likely, a response to drought signalling hormones resulting in the suppression of growth. For example, it will be interesting to investigate whether *irx9* dwarfing can be suppressed by an ABA-insensitive (ABI) mutant, such as *abi1* (Koornneef and van der Veen, 1980).

Indeed, some cell wall mutants have been reported as showing enhanced resistance to drought e.g. *irx14* (Keppler and Showalter, 2010) and *lew2* (a mutation in *CESA8* (Chen et al., 2005)), but also other abiotic stresses such as cold e.g. *esk1/tbl29* (a xylan acetyltransferase (Xin and Browse, 1998;Urbanowicz et al., 2014)). Furthermore, contrary to intuitive assumptions that modified walls will make plant cells more susceptible to pathogen entry, many cell wall mutants show increased resistance to pathogens e.g. (Manabe et al., 2011;Delgado-Cerezo et al., 2012), as has been reviewed recently (Miedes et al., 2014). When investigated in detail, it was demonstrated that the defence response genes were activated via an abscisic acid (ABA) signalling pathway (Hernandez-Blanco et al., 2007). Whilst many hypotheses have been proposed as to why this occurs, our data supports the possibility that the collapsed vessels “prime” the plants’ abiotic stress response, by simulating drought stress.

## 4 Conclusions

In this study, we utilised a collection of xylan mutants to establish whether the plant secondary cell wall has a cell wall integrity (SCWI) pathway whose members or consequences could be identified via transcriptomics. Despite using mutants with mild defects (*gux2*) through to mutants with severe defects (*irx9*), there was no transcriptional evidence of a SCWI pathway (Figure 2). This supports previously published biochemical data from secondary cell wall mutants, where, unlike in primary wall mutants, there was no detection of the compensatory deposition of polysaccharides (Ha et al., 2002;Cano-Delgado et al., 2003;Brown et al., 2007).

Transcriptomics does not necessarily reflect the dynamics of the cell, and therefore other approaches such as proteomics may be of use in the future to continue to search for the presence of a SCWI. Additionally, we have only investigated a single species in this study. It may be that perennial or monocotyledonous plants show altered responses to secondary cell wall changes (Tan et al., 2015).

The secondary cell wall is still a system about which we know little, despite advances over the last decade. In particular, mechanisms of *in muro* polysaccharide modification and salvage are poorly characterised. We also know little about how the characteristics of the different polymers are combined to produce the cell wall functionality. The identification of further cell wall mutants, along with the application of techniques which allow atomic resolution analysis will provide new insights into the mechanics of cell wall strength.

We believe that this data provides encouraging support for cell wall engineering approaches, especially for improved biomass properties, since there is not an intrinsic and sensitive regulatory system to overcome. We predict that as long as the plant is able to maintain form and function, e.g. transpiration flow (Petersen et al., 2012;Eudes et al., 2015;Vargas et al., 2016) by maintaining a functional secondary cell wall, it will be possible to manipulate the cell wall to produce biomaterials to suit different agricultural and biomanufacturing needs. Further work will be required to understand the contributions of interactions between individual wall components to form the assembled structure, to allow the predictive design of a functional secondary cell wall.

## 6 Author Contributions

JCM, NFB and PD designed the experiments and wrote the paper. NFB performed the experiments. NFB and JCM analysed the data.

## 8 Acknowledgments

The authors would like to thank Xiaolan Yu, who took the photograph of the mutants used in this study.

## 12 Supplemental Data

**Supplemental DataSet 1:** This excel file contains 3 worksheets. (1) All GCRMA normalized data (2) Examples of expression of some secondary cell wall synthesis genes in individual samples, to demonstrate that we harvested material actively transcribing the genes of interest (3) Extracted data for probeset 264493_at for IRX10 (At1g27440), showing the differential response of different oligonucleotide probes in the *irx10* mutant sample, depending on the position of the probe.

**Supplemental Dataset 2:** Lists of genes which are significantly up or down regulated in each mutant.

**Supplemental Dataset 3:** Extracted gene expression data for genes known to be involved in the synthesis or regulation of the following secondary cell wall biosynthetic processes: UDP-sugars, S-adenosylmethionine, acetyl Co-A, cellulose, mannan, xyloglucan, pectin, arabinogalactan proteins and transcription factors.

**Supplemental Figure 1:** PCR to confirm that all T-DNA lines used in this study were homozygous for the insertion of interest. g/g = amplification of DNA using gene-gene specific primers. g/i = amplification of DNA using gene-insert specific primers, where the insert specific primer is the left border of the T-DNA insertion. Blank = water was used in place of DNA in the PCR reaction.

**Supplemental Figure 2:** GO analysis of xylan mutants. (**A**) *irx9* (**B**) *irx10* (**C**) *irx14* (**D**) *gux1* (**E**) *gux2* (**F**) *gux1gux2*. The Classification Superviewer tool from the Bio-Analytic Resource for Plant Biology was used. Data for each category was normalised to the number of times it appeared in the GO analysis divided by the total number of genes assigned to that category.

